# Understanding the Causal Impact of Elevated Maternal Stress During Pregnancy: A Systematic Literature Review of Guinea Pig Models

**DOI:** 10.1101/2025.07.30.667555

**Authors:** Rebecca L. Wilson, Caitlin Messiah, Adetola F. Louis-Jacques

**Affiliations:** Center for Research in Perinatal Outcomes, University of Florida College of Medicine, Gainesville, Florida, USA 32610; Department of Obstetrics and Gynecology, University of Florida College of Medicine, Gainesville, Florida, USA 32610

**Keywords:** Maternal stress, pregnancy, guinea pig, developmental programming, review

## Abstract

Increased maternal stress during pregnancy has been linked to numerous adverse pregnancy outcomes, including preterm birth and impaired fetal development. While clinical studies have established strong associations between maternal stress and adverse pregnancy outcomes, understanding the biological mechanisms is crucial for developing effective preventive intervention. Rodent models have provided valuable insights, however physiological differences from humans limit translational relevance. This systematic review evaluated and synthesized experimental models of increased maternal stress during pregnancy in guinea pigs. Guinea pigs share key pregnancy characteristics with humans, including relevant placental structure, maternal hormonal recognition of pregnancy and precocial offspring development. We identified 43 studies using various experimental approaches categorized into three main types: behavioral models, drug/substance exposure models, and physiological challenge models. These studies demonstrated significant fetal and offspring effects, particularly sex-specific neurodevelopmental outcomes. However, maternal physiological adaptations were often superficially characterized, focusing primarily on weight gain and occasional stress hormone measurements. Most studies began interventions during established pregnancy, potentially missing critical pre-pregnancy and early pregnancy periods. Overall, this review highlights the need for future studies to consider pre-conception interventions to better model human conditions and more comprehensively examine maternal adaptations.

## Introduction

Increased maternal stress, including psychological stress (example, anxiety or depression) and physical stressors (example, malnutrition or infection), during pregnancy has been linked to numerous adverse pregnancy outcomes in humans. Clinical studies have demonstrated associations between maternal stress and increased risk of preterm birth (Dole et al., 2003; Copper et al., 1996), low birth weight (Wadhwa et al., 2004) and preeclampsia (Zhang et al., 2013). Additionally, the impact of high maternal stress during pregnancy has been shown to have extensive effects on fetus and offspring health and development [1, 2].

The biological mechanisms underlying the associations between high maternal stress and adverse pregnancy outcome are complex but likely involve a number of key physiological mechanisms. Activation of the maternal hypothalamic-pituitary-adrenal (HPA) axis, resulting in elevated cortisol levels can affect placental function and fetal development [3, 4]. Maternal stress can trigger inflammatory responses and alter immune function, potentially compromising pregnancy maintenance [5]. Additionally, the timing of stress exposure appears particularly important, with different gestational periods showing varying susceptibility to stress-induced complications [6, 7]. While human studies have provided valuable insights into these relationships, animal models are essential for understanding the causal relationships between increased maternal stress, pregnancy outcome and long-term offspring health.

Rodent models are a cost-effective option for studying causal relationships and have been valuable in investigating potential mechanisms linking maternal stress to adverse pregnancy outcomes. Most consistent findings across rodent models are those relating to fetal growth and placental function [8–10]. Outcomes often reported include reduced litter size and litter weight and activation of the maternal HPA-axis that is associated with reduced circulating progesterone and alterations in placenta structure and functions. However, while human studies consistently show associations between maternal stress and increased risk of preterm birth [11], rodent models have produced variable results regarding gestational length [12]. In those that did report early delivery, gestational length was only minimally premature [13, 14]. The variability in pregnancy outcomes might reflect differences in stress protocols, timing of exposure, or species-specific responses to maternal stress [15].

The guinea pig (Cavia porcellus) has emerged as a particularly valuable model species for studying maternal-fetal interactions due to its similarities to human pregnancy, including comparable placental structure, relatively long gestation period, and advanced fetal development [16–18]. Guinea pigs, unlike rats and mice, exhibit similar hormonal profiles to humans during pregnancy, particularly in their production and regulation of progesterone, estrogen, and stress hormones [19, 20]. The maternal hypothalamic-pituitary-adrenal (HPA) axis undergoes comparable adaptations during pregnancy, including increased basal cortisol levels and altered stress responsiveness [21]. Importantly, like humans, but unlike most rodents, guinea pigs primarily produce cortisol as their main glucocorticoid, rather than corticosterone [22]. This is significant because cortisol plays a crucial role in maternal metabolic adaptation to pregnancy and the timing of parturition in both species [23, 24]. Additionally, the guinea pig’s relatively long gestation period (∼65-70 days) allows for detailed study of temporal changes in maternal physiology across pregnancy, more closely matching the timeline of human gestational adaptation than shorter-gestation rodent models.

Over the past two decades, researchers have developed various experimental approaches to investigate the causal connections between increased prenatal maternal stress, pregnancy outcome and offspring developmental outcomes using guinea pig models (see references summarized in this review). These studies have encompassed a wide range of interventions, from psychological stressors to physiological challenges, providing insights into both the immediate and long-term consequences of prenatal stress exposure, particularly in offspring. The diversity of experimental approaches means multiple aspects of fetal and offspring development, including neurodevelopmental outcomes, cardiovascular function, and metabolic programming have been examined. This systematic review aimed to evaluate and synthesize the available literature on maternal stress during pregnancy using guinea pig models, focusing on experimental manipulations published between 2000 and present. We specifically examine the types of interventions employed, the timing of these interventions during pregnancy, and their effects on both maternal and offspring outcomes. This analysis will provide a critical overview of current knowledge in the field and identify gaps that require further investigation.

## Materials and Methods

### Eligibility Criteria

Studies included only those utilizing guinea pigs as a model species and focused on maternal experimental manipulation during pregnancy. There were no restrictions imposed on type of experimental manipulation but was specifically targeted to increased maternal stress, either psychologically or physiologically, during pregnancy. Given the heterogeneity of the observational strategies, a meta-analysis was not possible.

### Information Sources and Search

The search strategy and procedure was guided by the PRISMA statement [25]. Potential studies were located through electronic databases (Web of Science, Academic Search Premier and Scopus), as well as manual searches of references in review articles and relevant articles known by the authors. Limits included full text articles written in English and published in academic journals between 2000-present. The last search was performed in June 2025. Search terms and MeSH headings in the title, abstract, and index terms, were initially identified in Scopus and subsequent key words were used for the remaining databases (Appendix A). The search included the following: Maternal Stress or Stress; Pregnancy or Prenatal; Guinea Pig or Guinea Pigs.

### Data Collection

An independent search of the literature was performed in April 2025 and again in July 2025. Titles and abstracts were examined independently by two of the authors who documented reasons for excluding full text articles. Any differences between the two reviewers were clarified; a third reviewer resolved any disagreements. If an article appeared in duplicate from two or three of the databases, only the search containing the most relevant and useful information was included. For each eligible study, the following data was extracted: author, year; aim/hypothesis; study design; study outcome and preterm birth phenotype.

## Results

Figure 1 outlines the literature search and selection of studies. We identified 172 citations (10 duplicates) after searching Web of Science, Academic Search Premier and Scopus databases. A further 16 were added by authors after handsearching. After screening the title and abstract, 55 full text papers were read. Overall, 43 studies met the inclusion criteria and were included in the review (Table 1).

**Figure 1.**
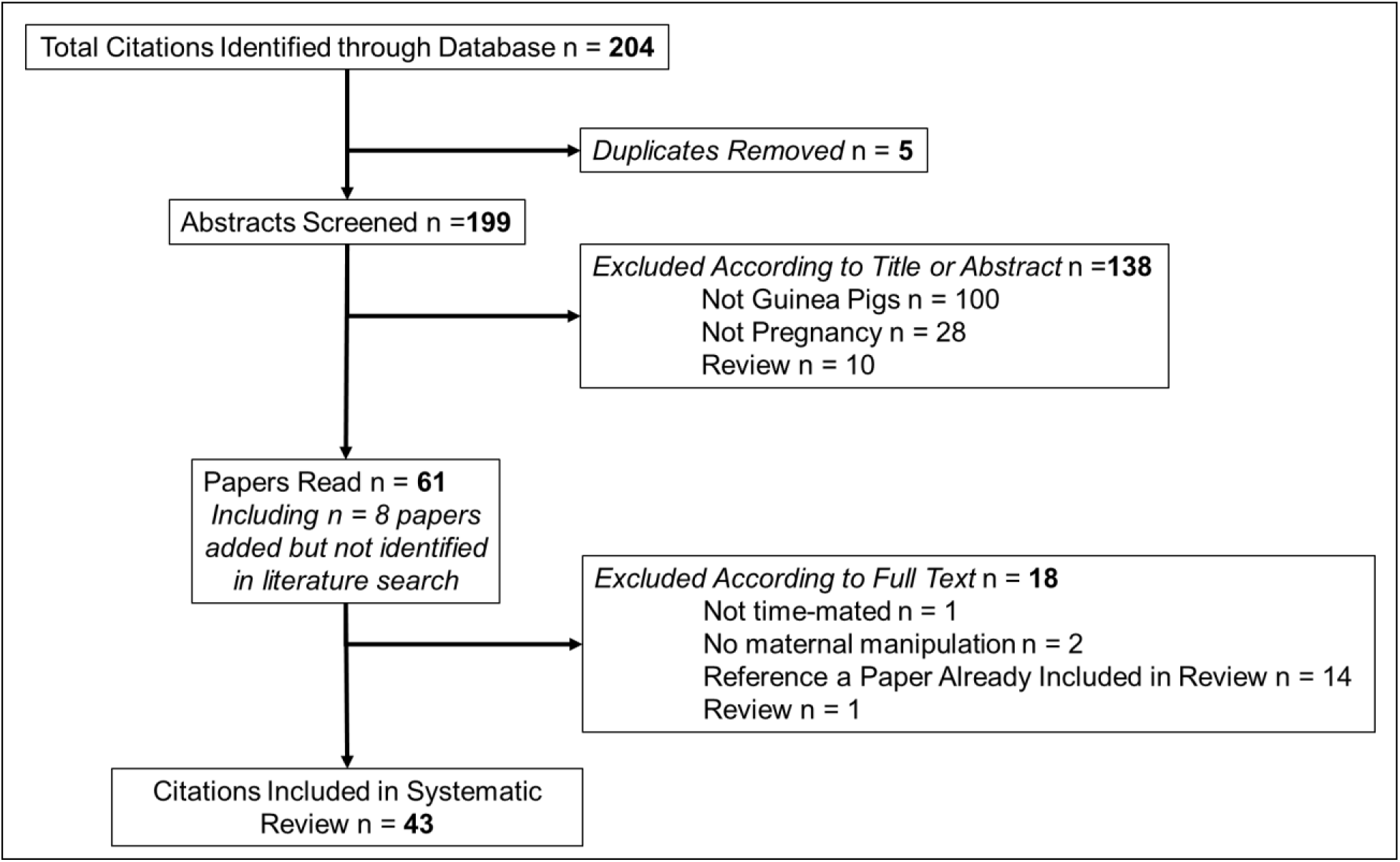
Flow diagram of the search strategy used in this review including the relevant number of papers at each point.

**Table 1.**
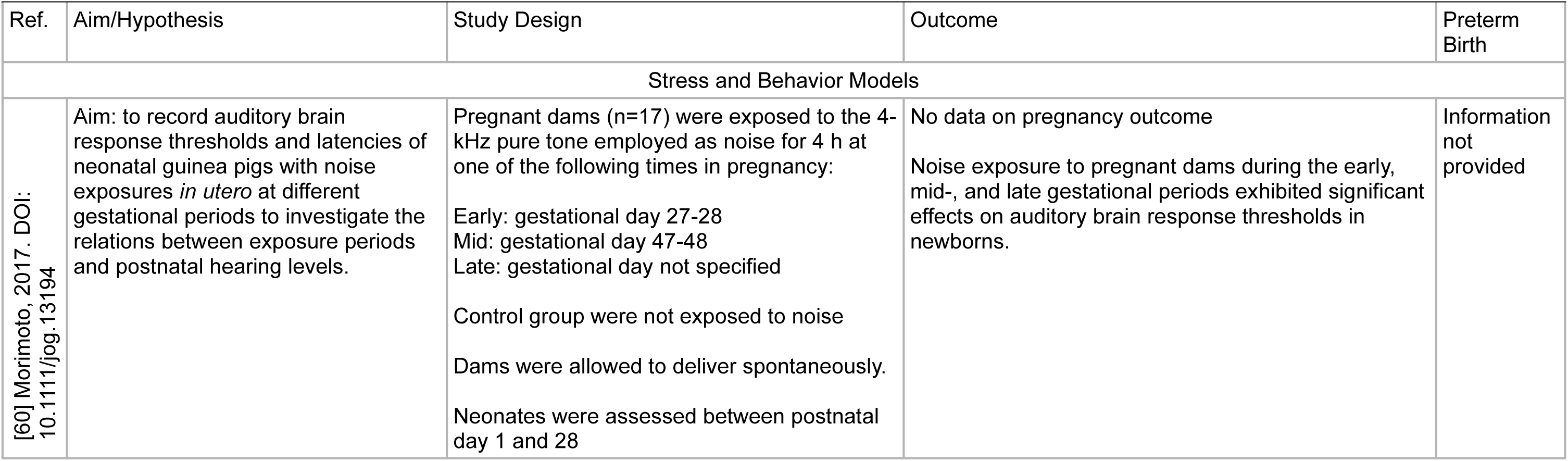
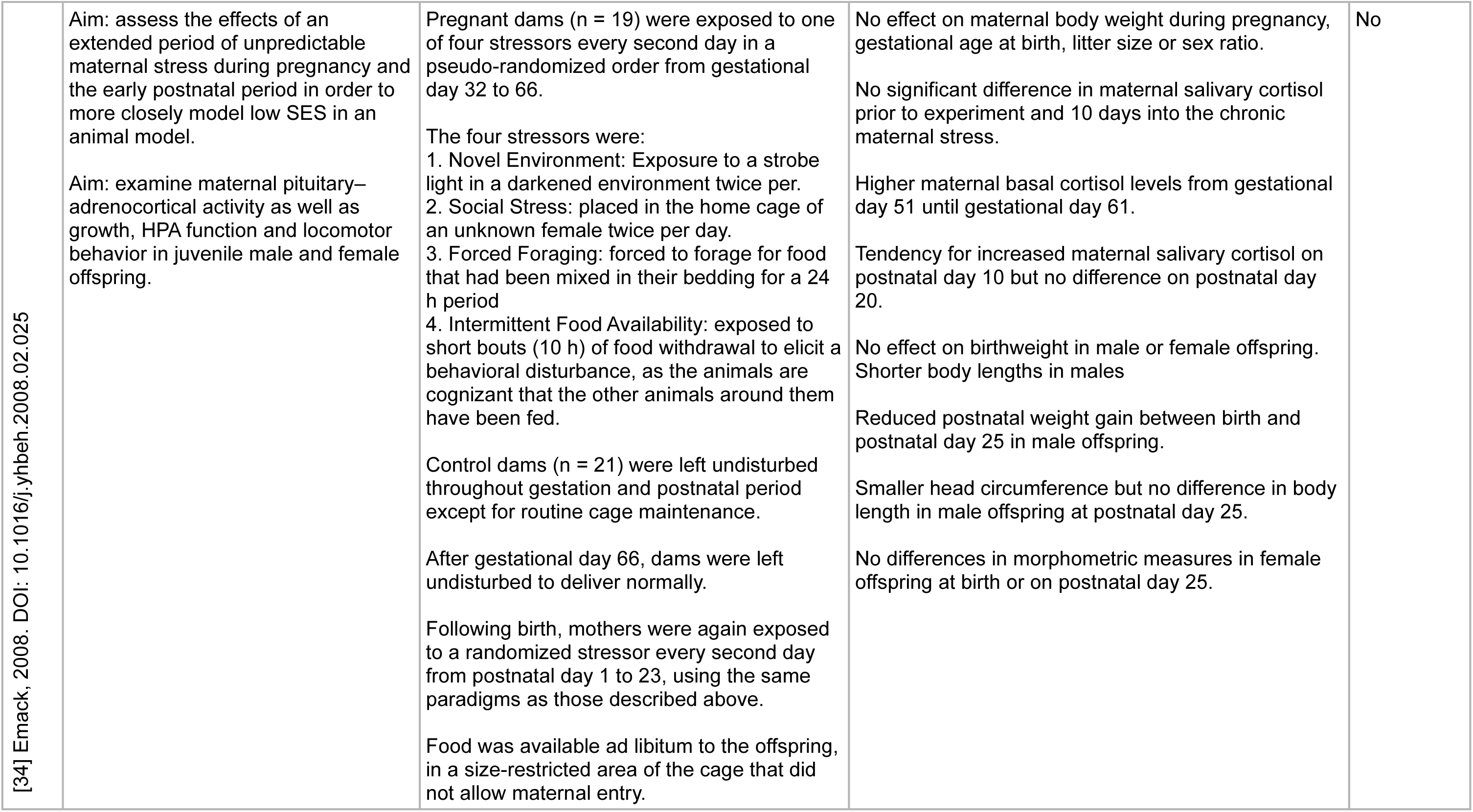
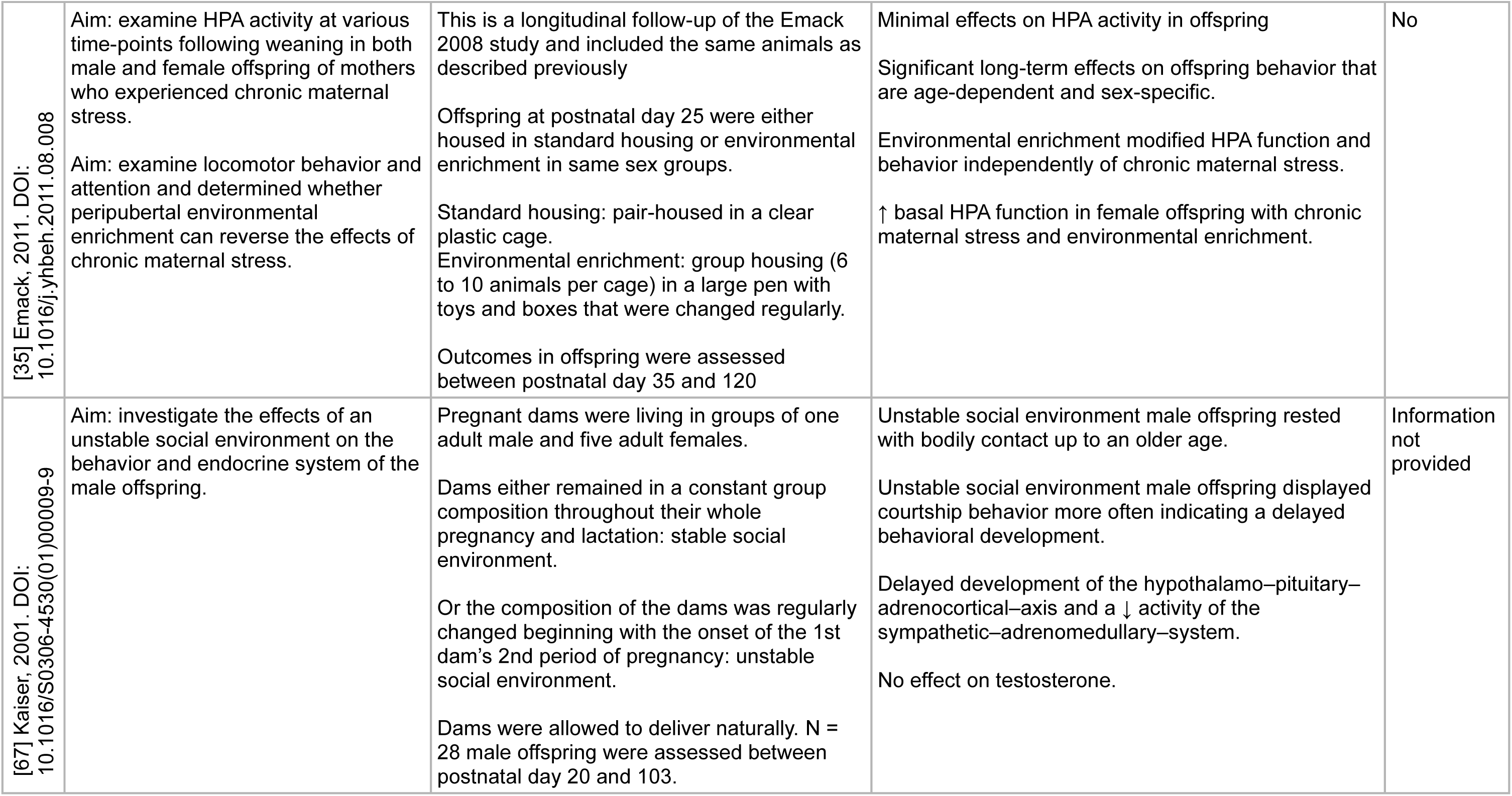
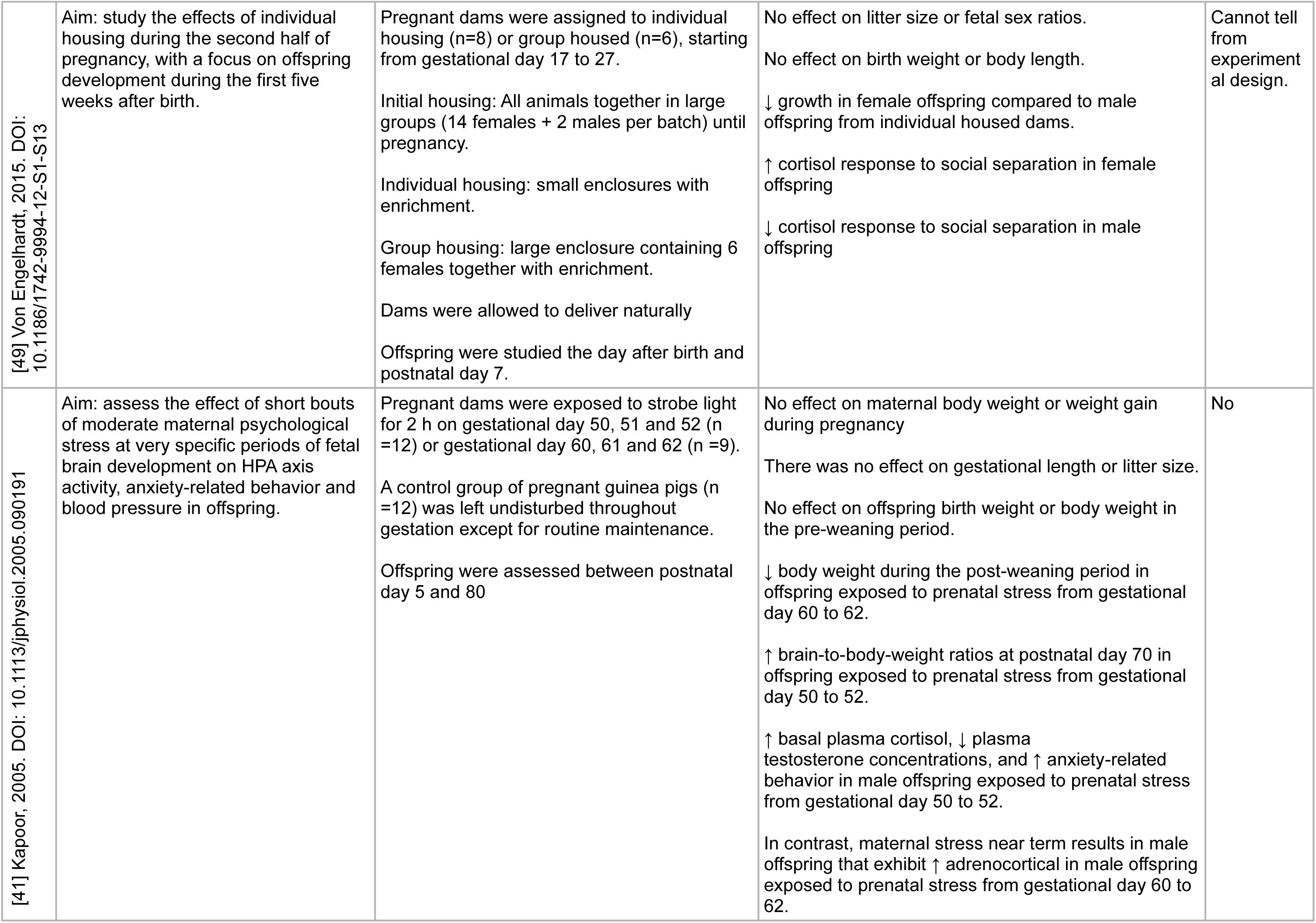
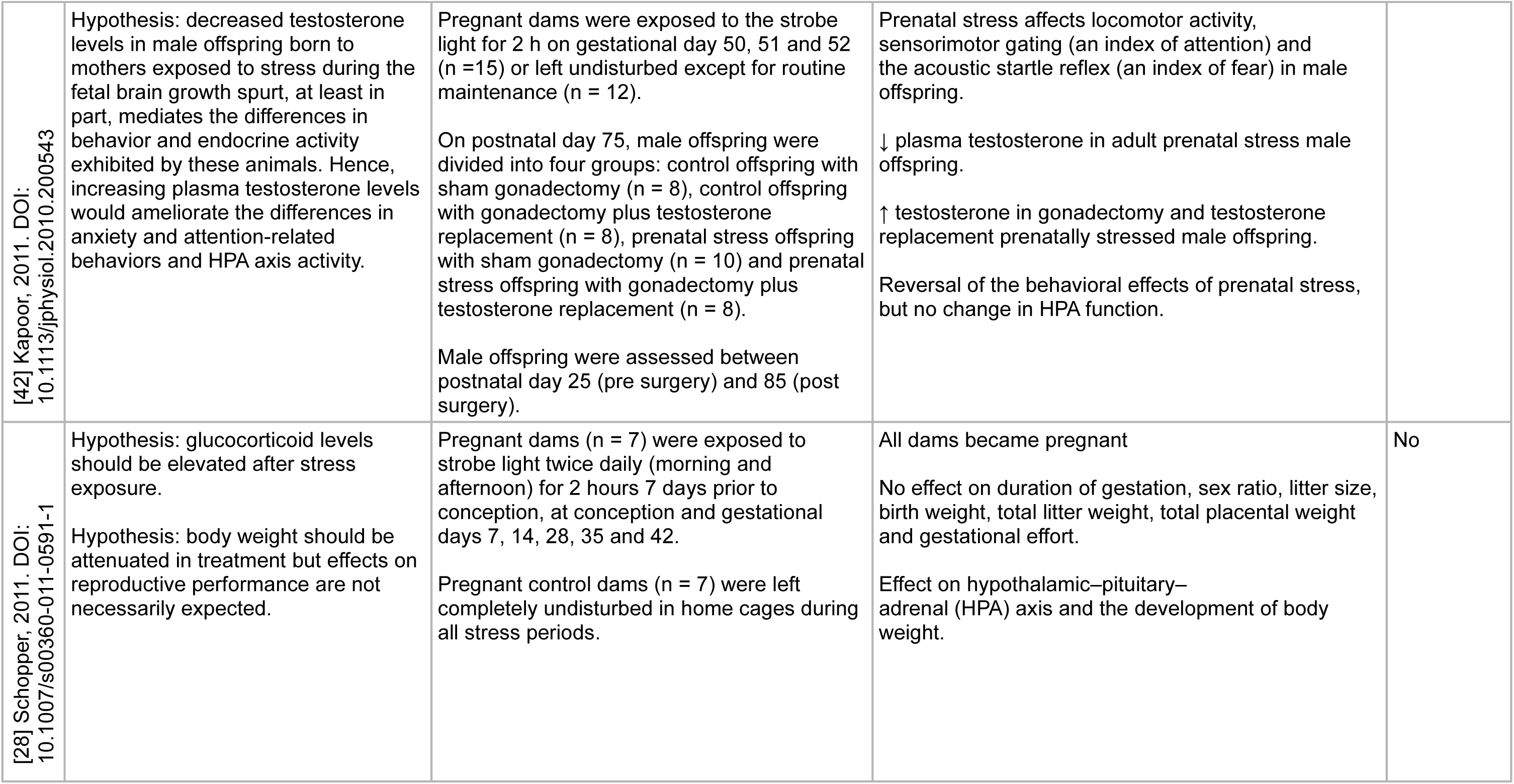
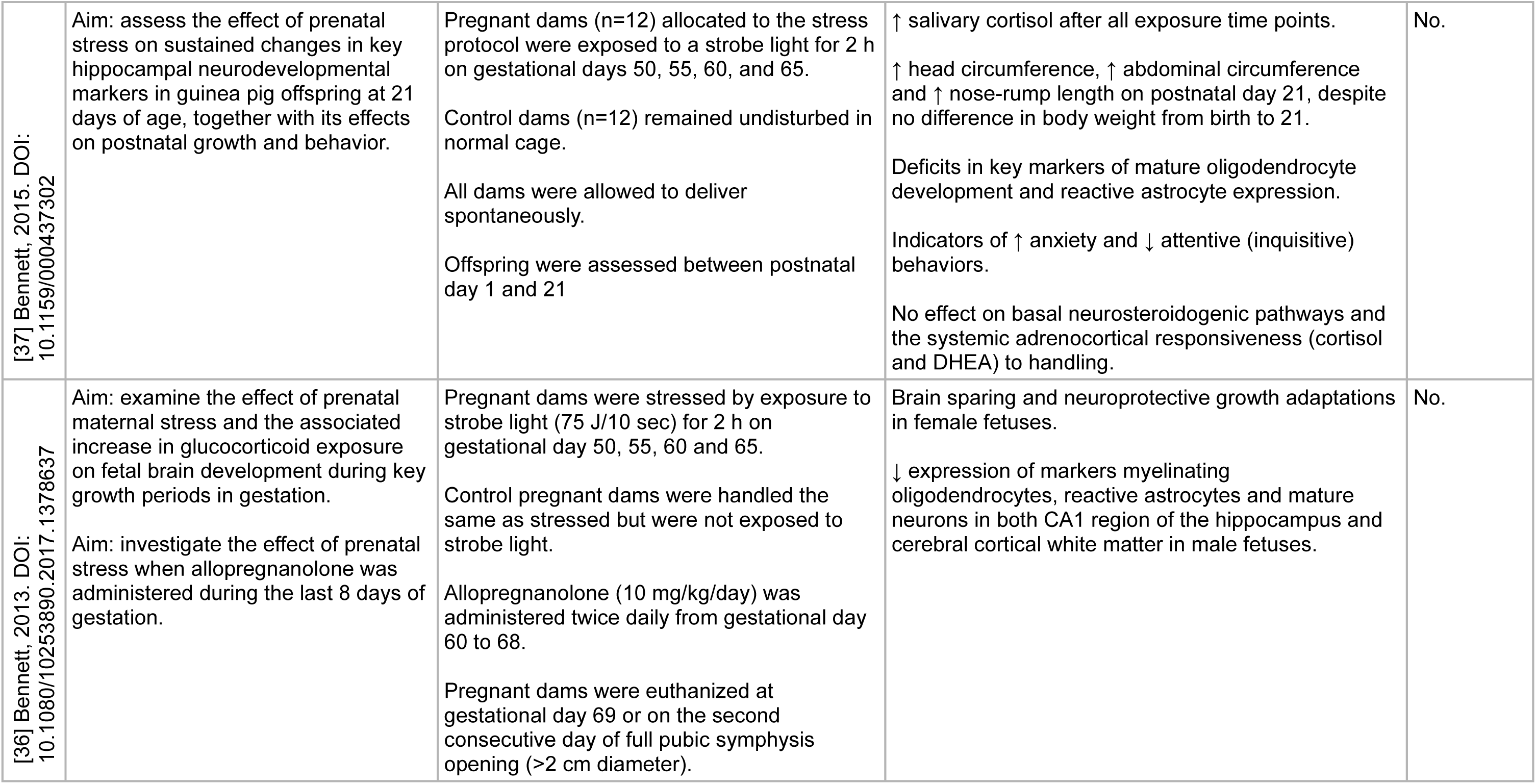
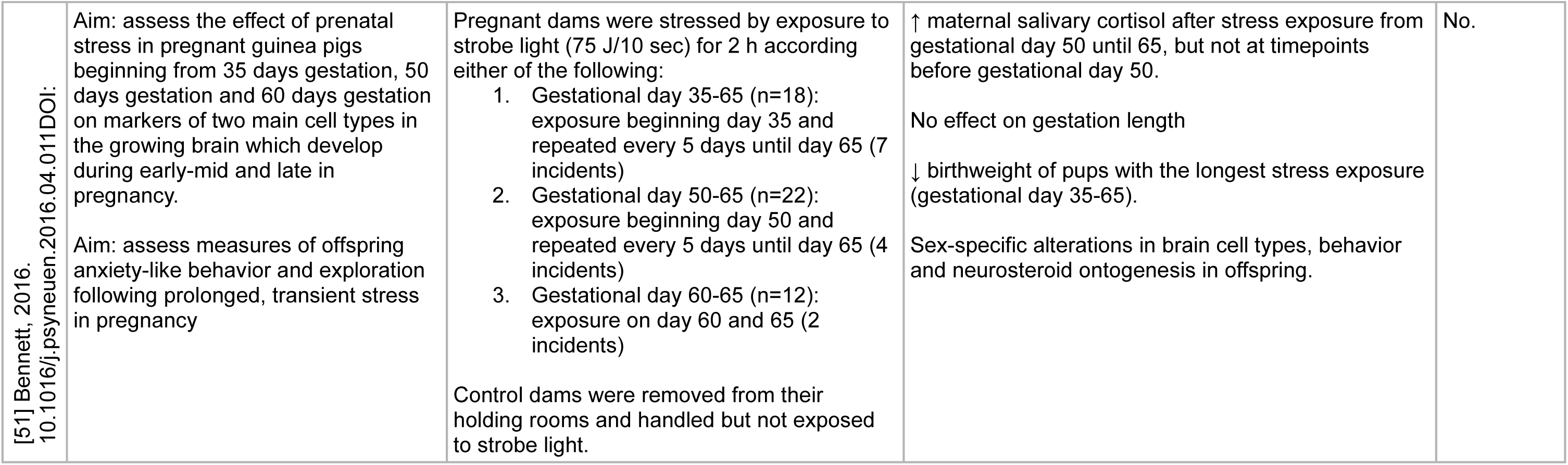
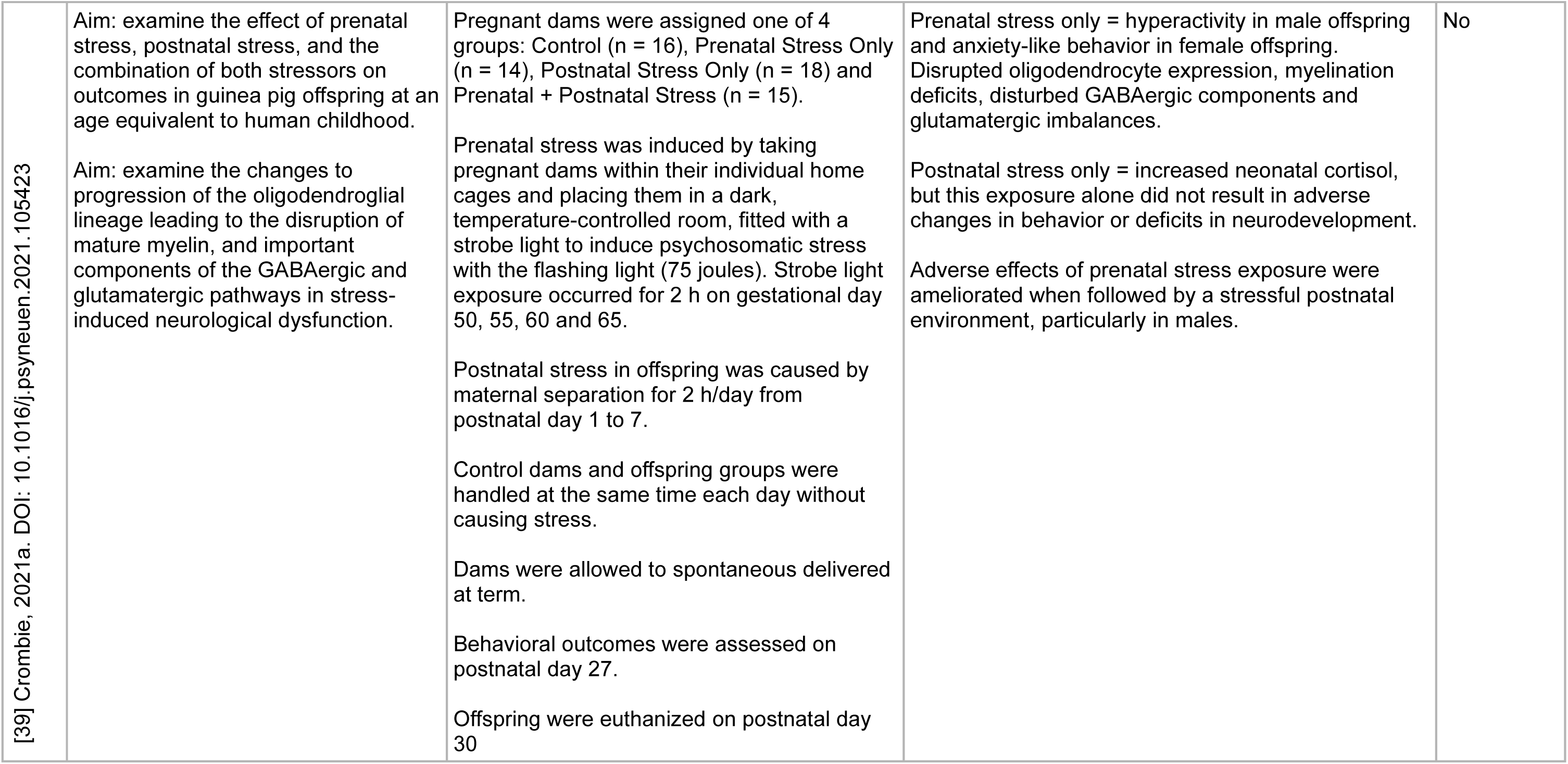
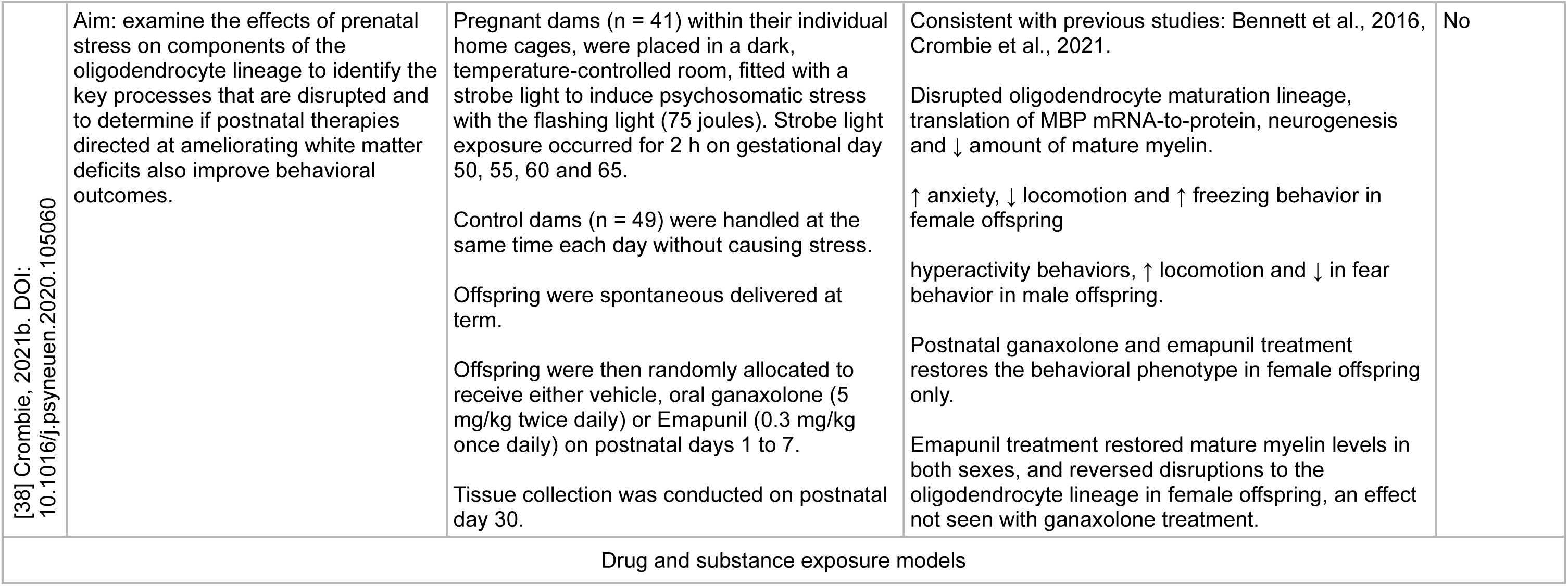
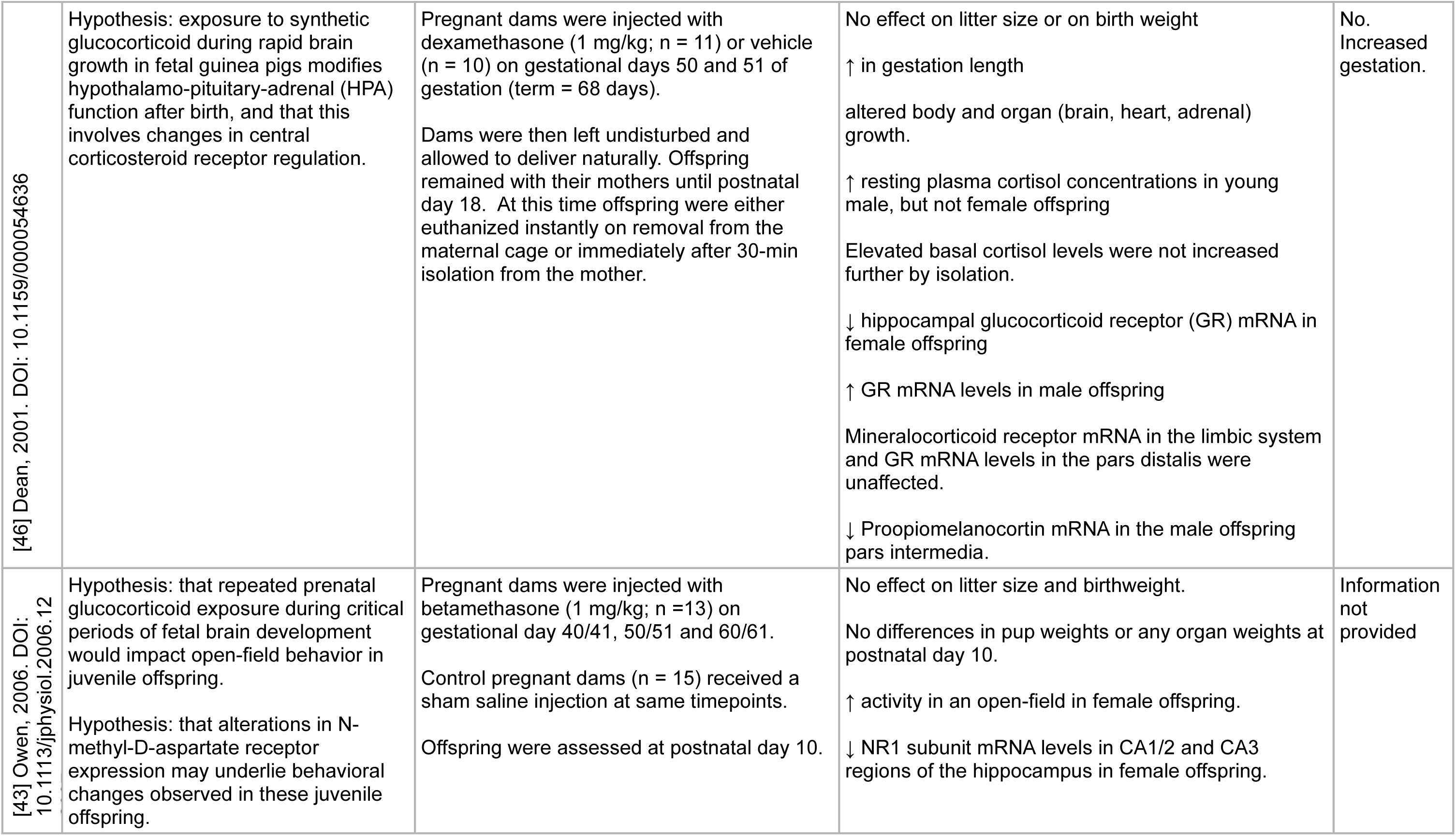
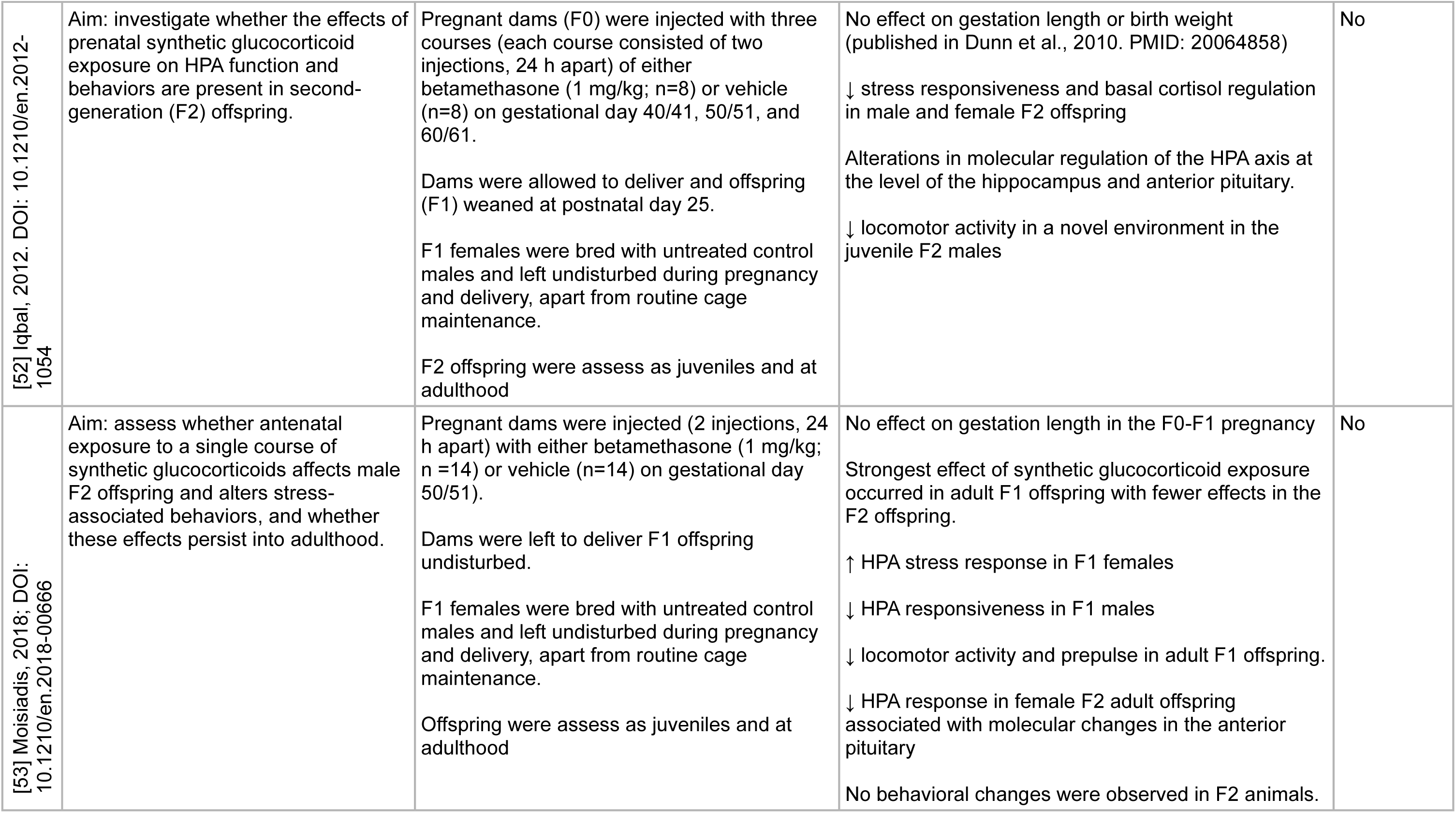
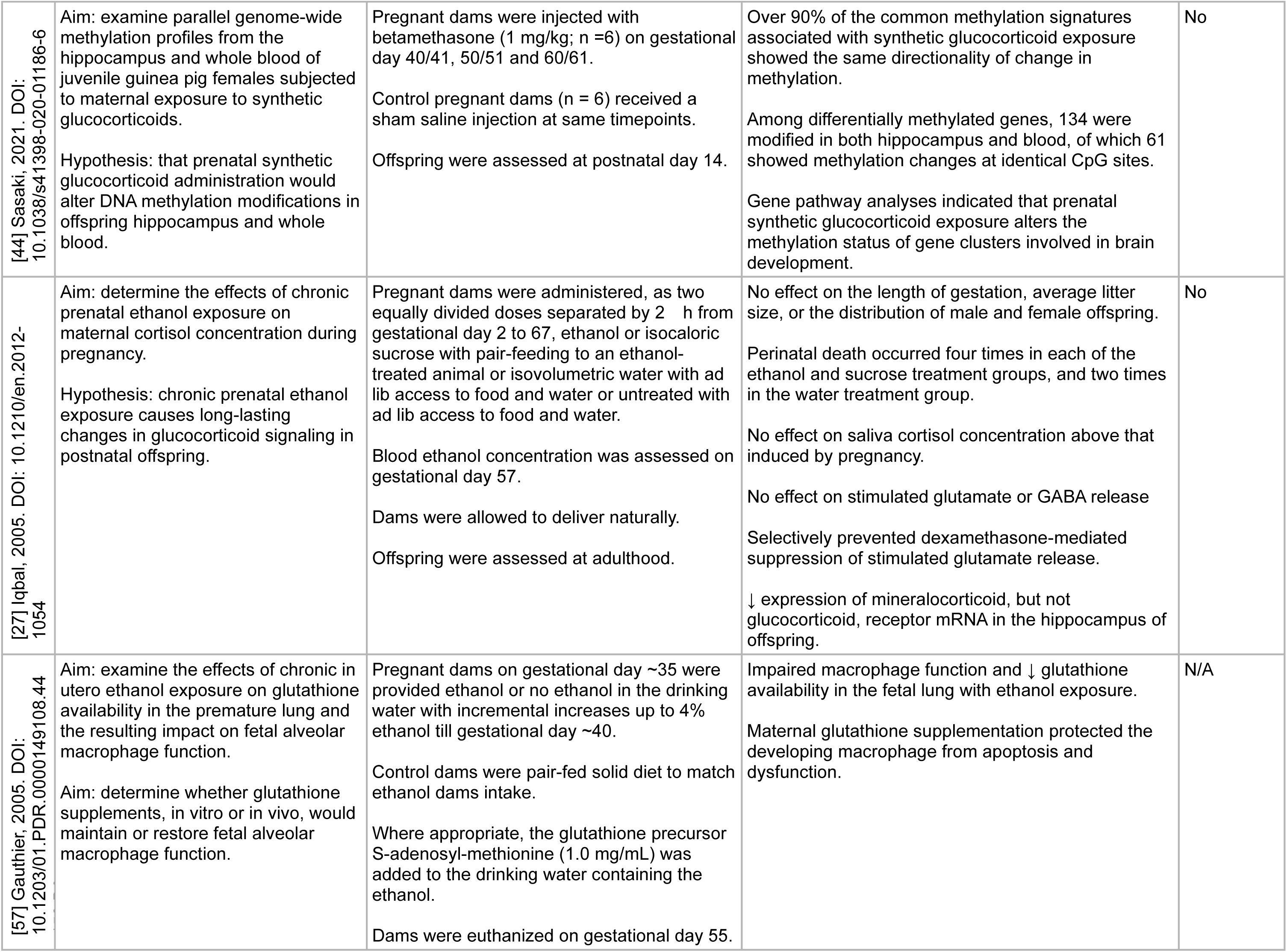
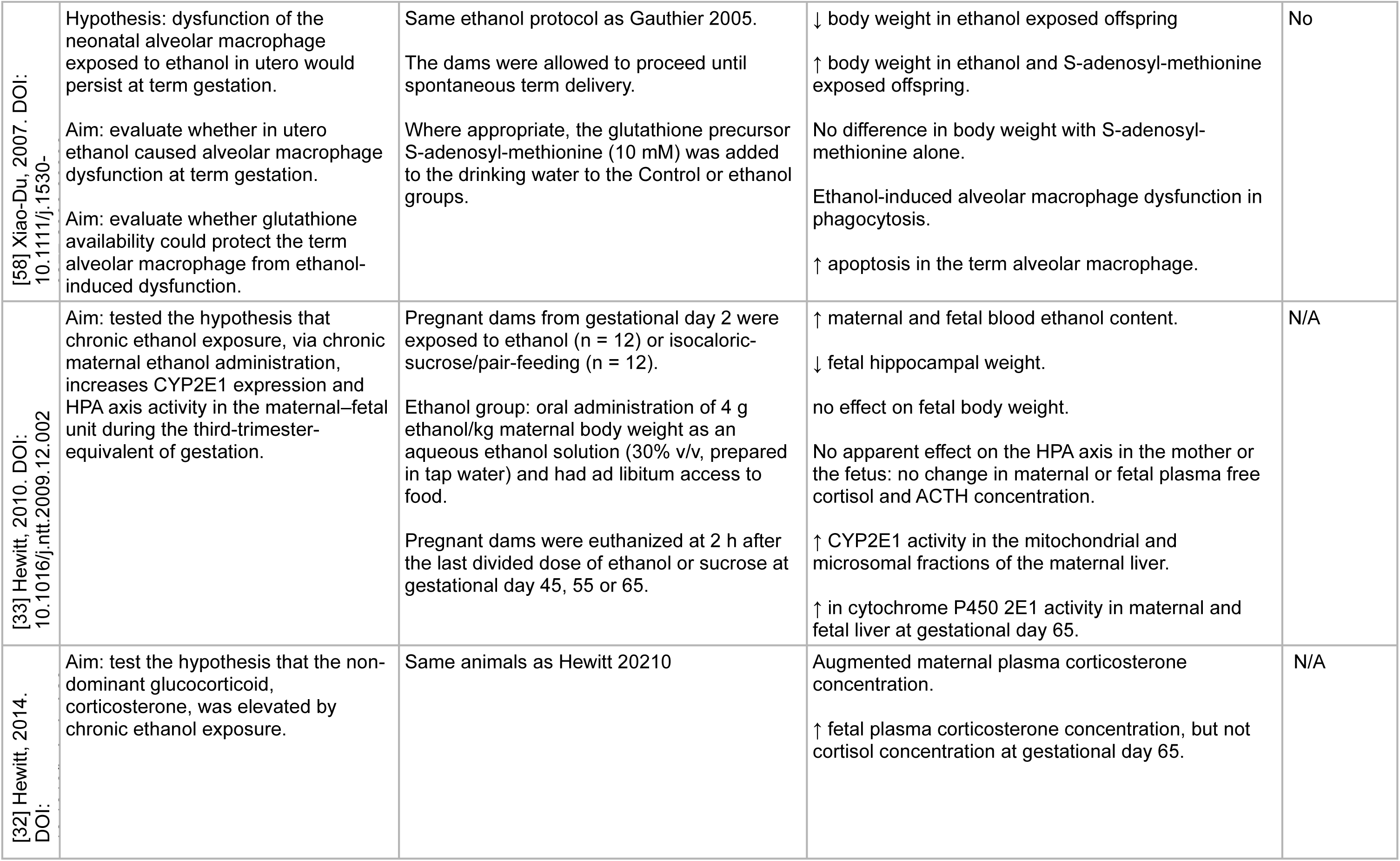
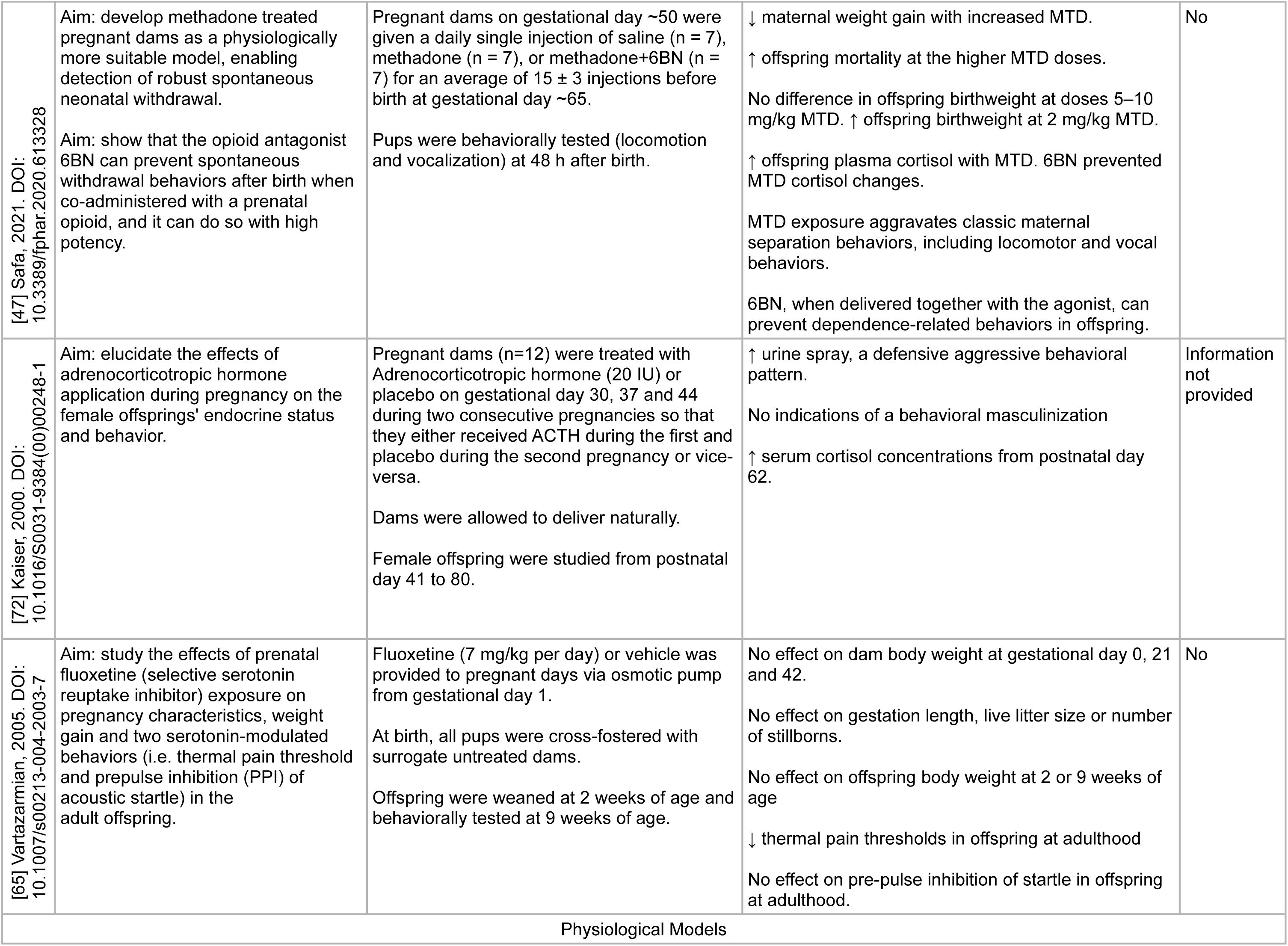
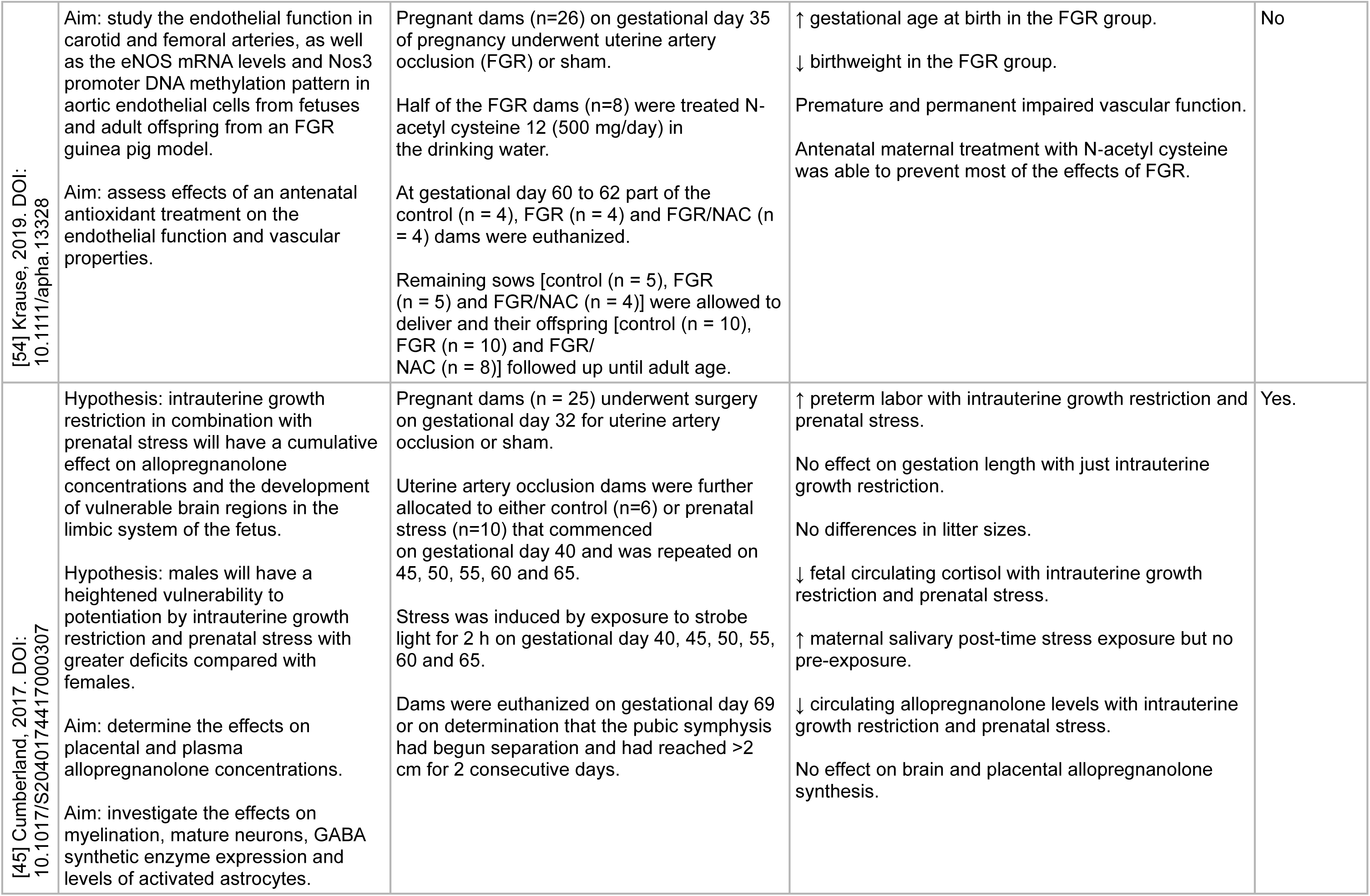
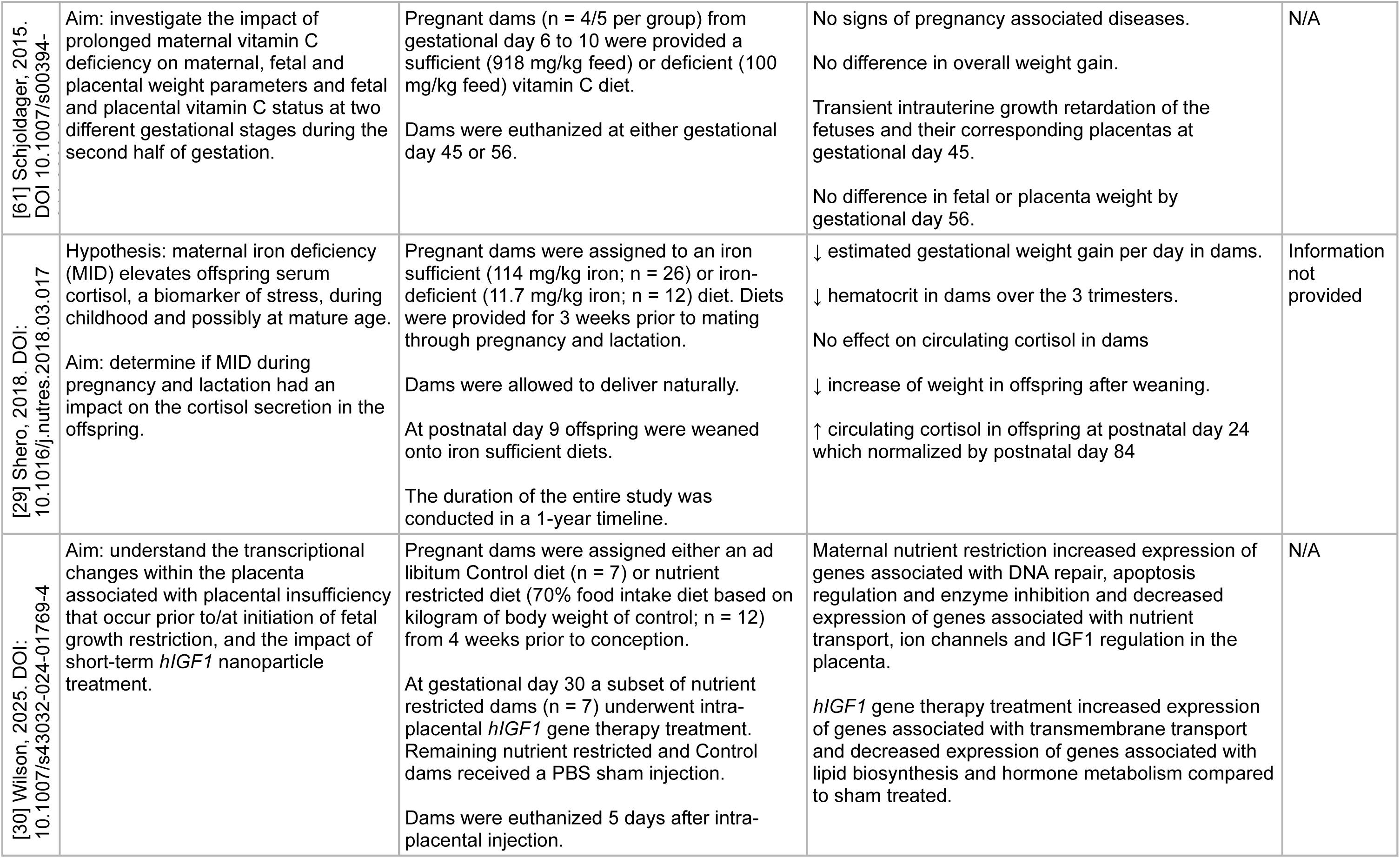
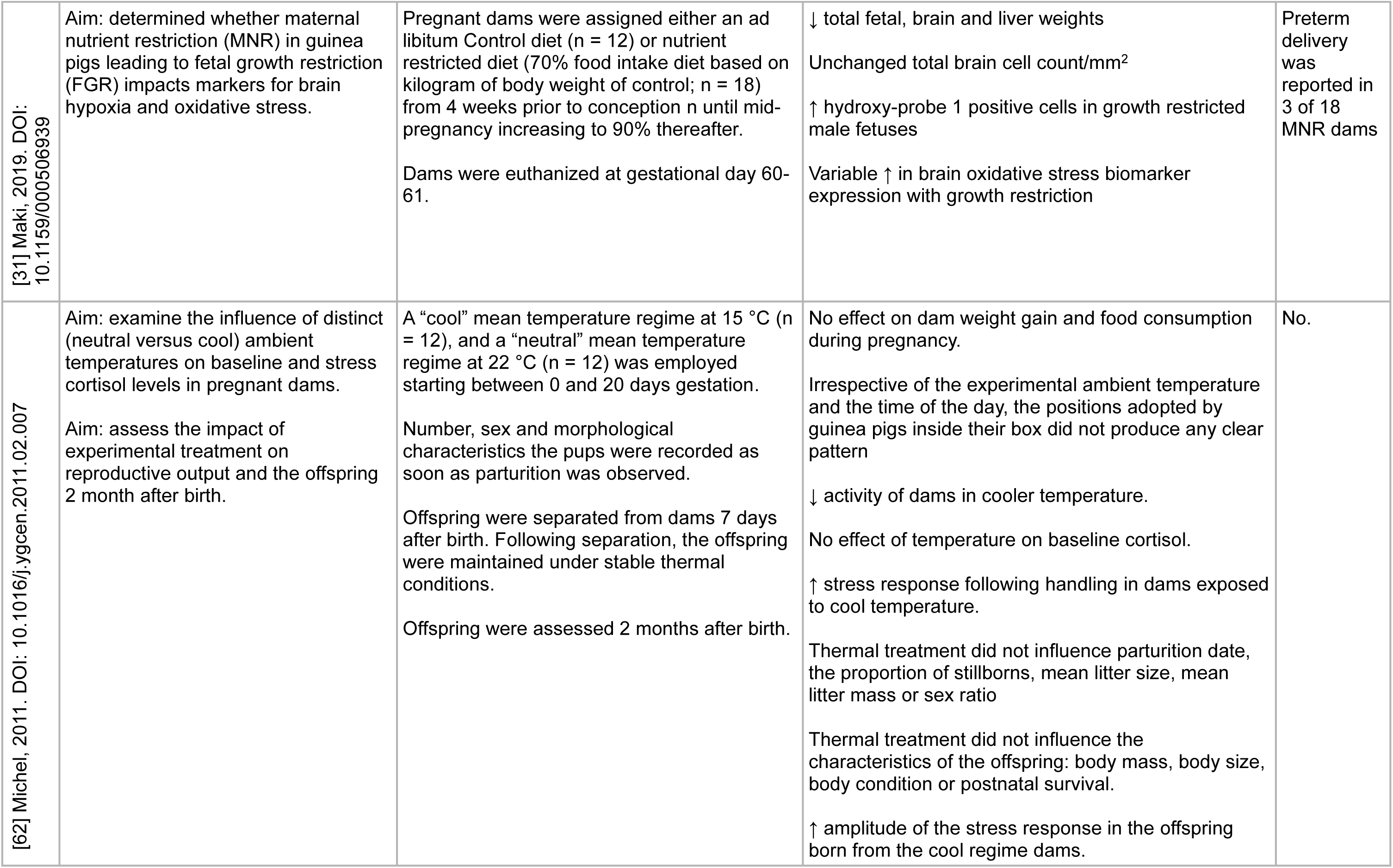
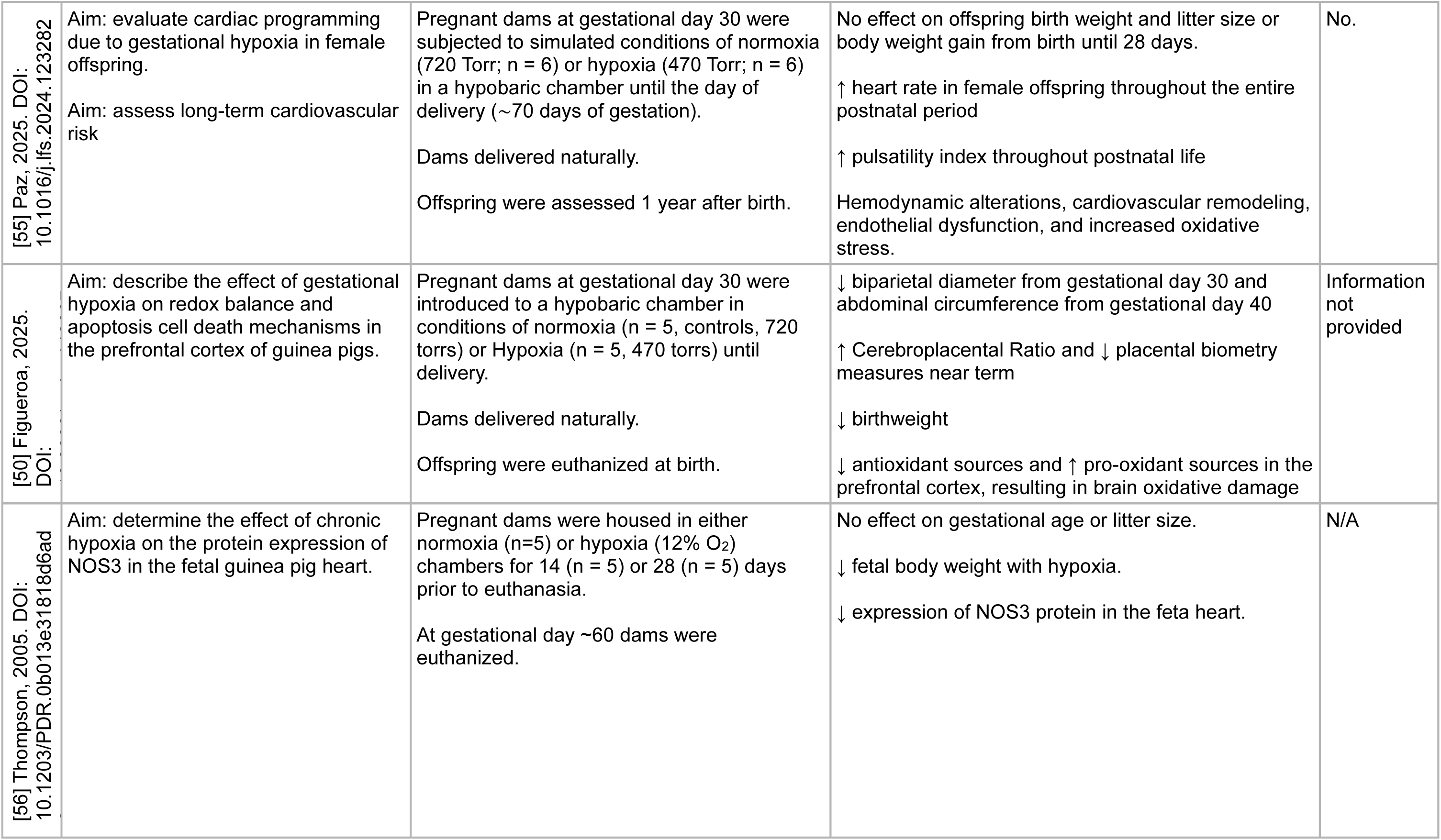
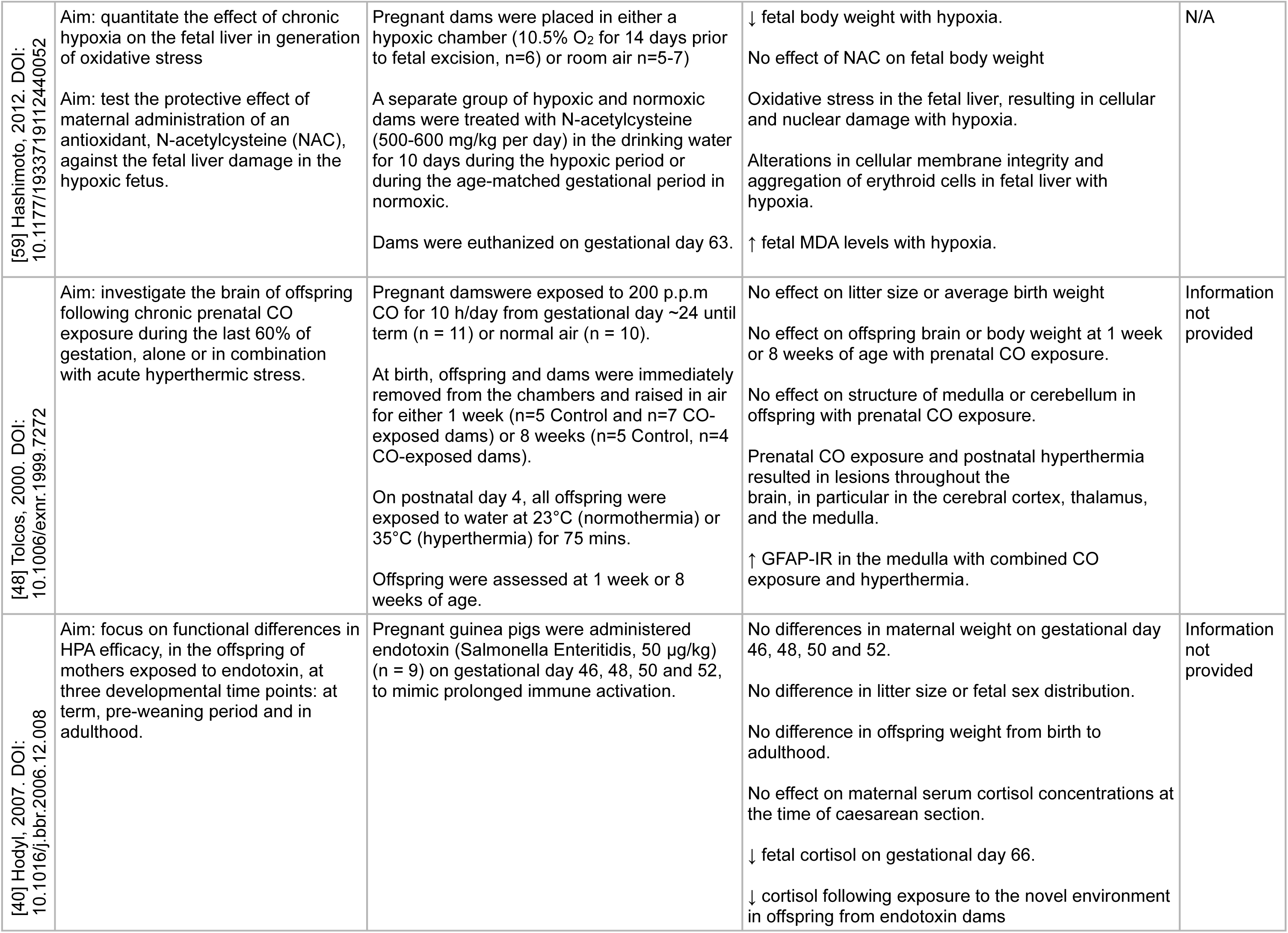
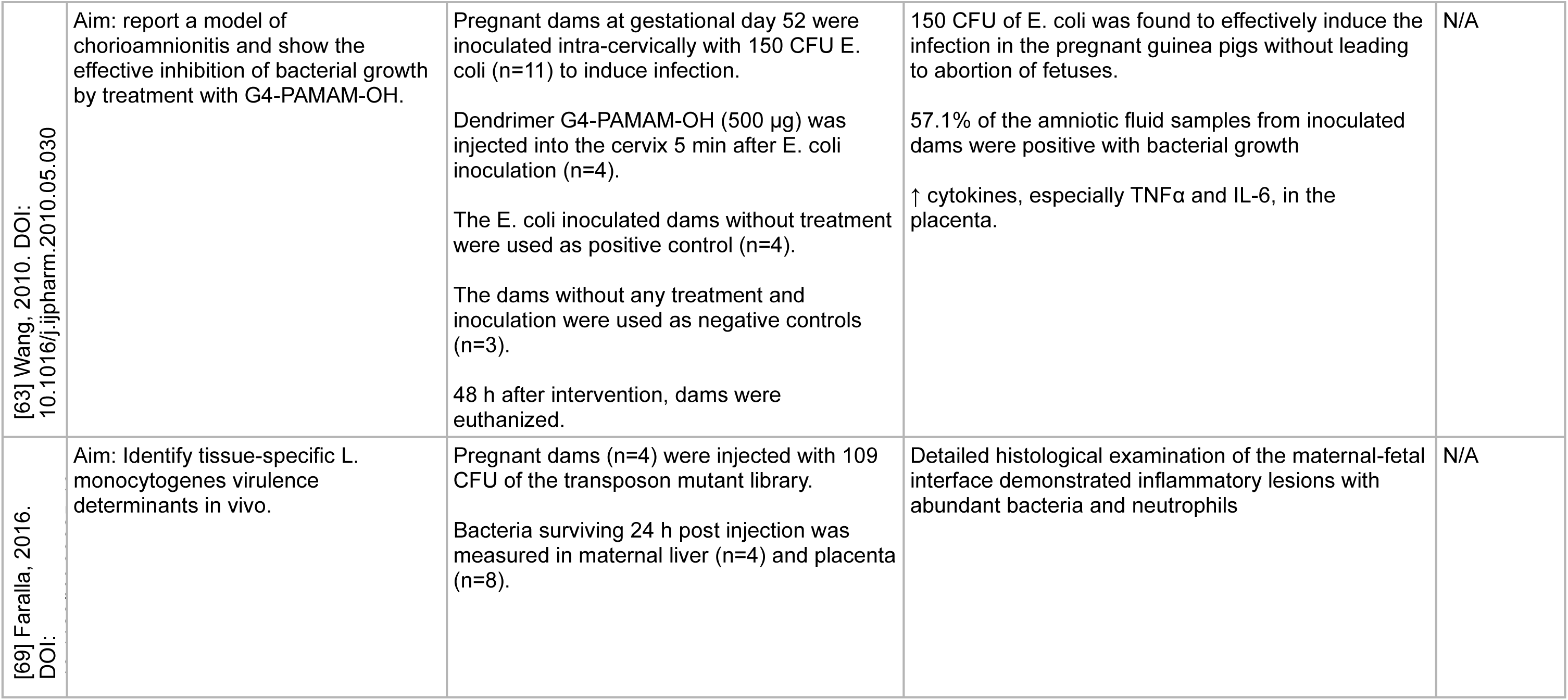
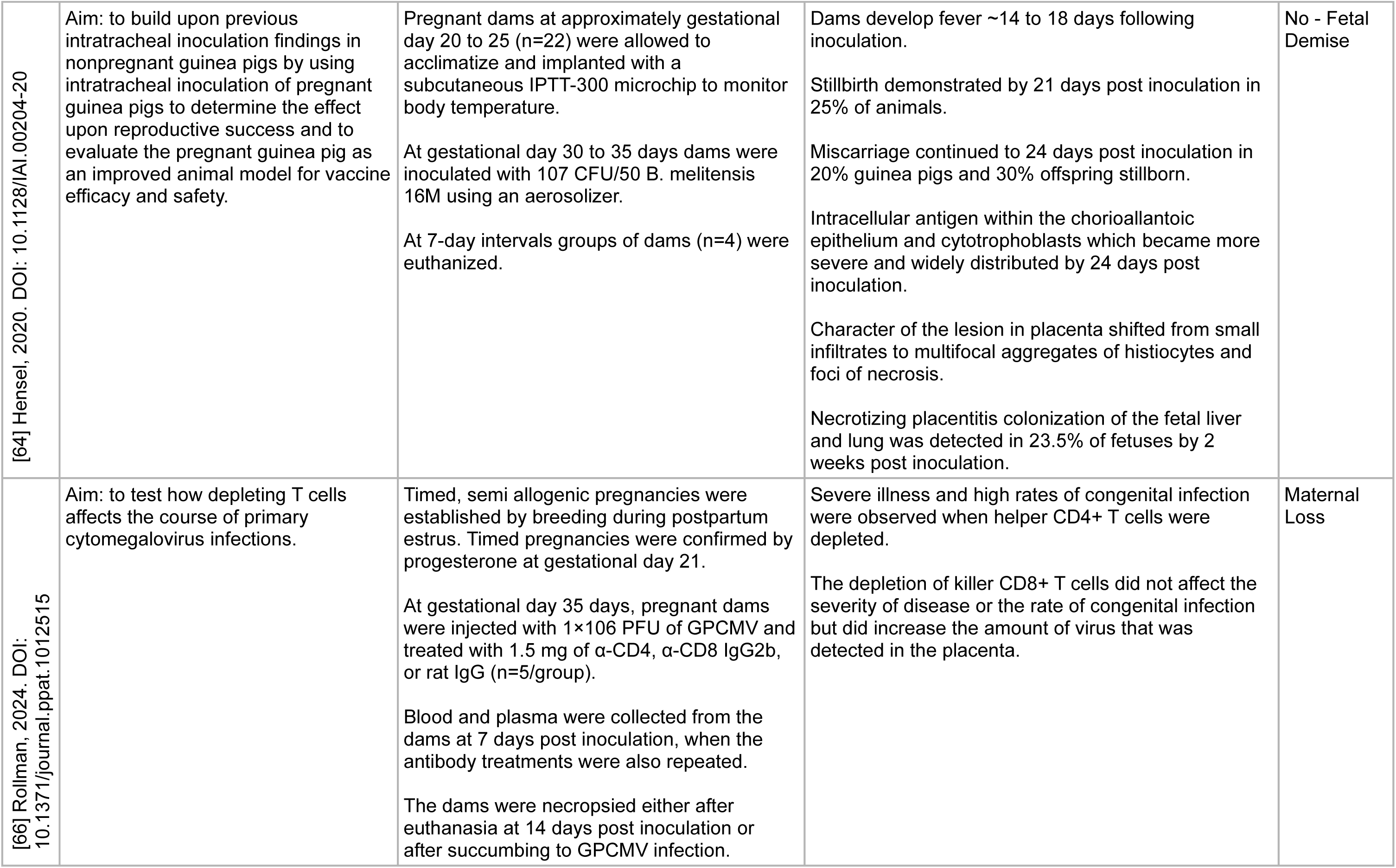
Summary of studies systematically reviewed.

Most studies utilized timed pregnant guinea pigs with interventions occurring at specific gestational ages, typically between days 45-65 of pregnancy (term 65-70 days) [26]. One study began experimental manipulations from gestational day 1 [27] while only four studies [28–31] utilized a pre-conception experimental manipulation. The experimental designs generally included relatively small sample sizes, ranging from 4-16 animals per group, with most studies examining fetal and/or offspring outcomes. The majority of studies aimed to assess near-term fetal brain development [32–35] or neurodevelopmental outcomes in offspring [27, 36–53]. Additional aims included understanding fetal and/or offspring vascular function [54–56], fetal and/or neonatal lung development and function [57, 58], fetal liver damage [59], offspring auditory outcomes [60] and maternal impacts [61, 62].

Regarding pregnancy outcomes, most interventions did not significantly affect gestational length, though several models showed reduced fetal/offspring weights [31, 33, 50, 54, 56, 58, 59]. Fetal survival was often demonstrated except in cases of severe interventions such as infection models [63, 64]. Impacts to offspring were routinely sex-specific in nature [31, 34–36, 38, 39, 41–43, 46, 49]. For example, Crombie et al., [39] found that following maternal psychosomatic stress via strobe light in pregnancy, female offspring showed increased anxiety, decreased locomotion and increased freezing while male offspring showed hyperactivity behavior, increased locomotion and decreased fear behavior. When reported, maternal responses to experimental manipulations were generally well-tolerated with dams maintaining appropriate weight gain throughout pregnancy [34, 40, 41, 61, 62, 65] except with iron deficiency [29] and methadone treatment [47] where maternal weight gain was reduced. Increased maternal cortisol and stress responses were observed with strobe light [37, 45] and combined environmental stressors [34] but not with hyperthermia [62], chronic ethanol exposure [27, 33], iron deficiency [29] or endotoxin administration [40]. Adverse maternal outcomes were relatively rare, primarily occurring in infection models [64, 66].

The experimental manipulations to increase maternal stress used in these studies can be categorized into three main types. Stress and behavioral models [34–39, 41, 42, 49, 51, 60, 67]: maternal stress induced via strobe light exposure, maternal separation protocols, novel environment exposure, and social stress paradigms. Drug and substance exposure models: betamethasone or dexamethasone administration [43, 44, 46, 52, 53], ethanol exposure [27, 32, 33, 57, 58], methadone [47] and pharmacological intervention [65, 67]. Physiological models: uterine artery occlusion for growth restriction [45, 54], maternal nutrition manipulation [29–31, 61, 68], hypoxia exposure [50, 55, 56, 59], carbon monoxide [48], temperature [62] and infection/inflammation challenges [40, 63, 64, 66, 69].

### Stress and behavioral models

Stress and behavioral models in pregnant guinea pigs have been particularly well-characterized, with several research groups establishing reproducible protocols that demonstrate clear offspring effects. The most commonly used protocol involved exposure to strobe light stress, typically administered for 2-hour periods during specific gestational windows [36, 41]. This model induced a reliable maternal stress response, evidenced by elevated salivary cortisol levels, without causing pregnancy loss or significantly altering gestational length [37]. The timing of stress exposure appears critical, with different windows of exposure producing distinct offspring phenotypes, particularly in behavioral and neuroendocrine outcomes [41, 51]. The physiological basis for these behavioral changes has been extensively investigated, with studies revealing alterations in offspring HPA axis function [28, 37] and modifications in neurotransmitter systems [43].

More complex stress paradigms have also been developed, including chronic unpredictable stress protocols that better mirror human psychological stress [34, 35] and social environment manipulations [28, 49, 67]. [34] demonstrated that combining multiple stressors, including novel environment exposure, social stress, and intermittent food availability, created a more comprehensive model of chronic maternal stress. This approach showed that maternal HPA axis activation persisted throughout gestation, with increased basal cortisol levels evident by mid-gestation. The offspring effects of these various stress protocols were sex-specific, with males generally displaying increased anxiety-like behaviors and hyperactivity, while females showed more subtle behavioral alterations [39, 42, 49]. Importantly, these behavioral changes persisted into juvenile periods, suggesting permanent programming effects [37]. Some studies have attempted to ameliorate these stress-induced changes through postnatal interventions or environmental enrichment, though with varying success [35, 38].

### Drug and substance exposure models

Drug and substance exposure models primarily focused on glucocorticoids and ethanol. Betamethasone and dexamethasone were studied because of their use to accelerate fetal lung maturation in cases of premature delivery [70, 71]. Early work by Dean et al., [46] showed that dexamethasone exposure during rapid brain growth altered offspring HPA function in a sex-specific manner, with increased resting cortisol in males but not females. Multiple studies examined both single and repeated courses of betamethasone, also administered at critical windows of brain development [43, 44]. Studies showed that maternal betamethasone administration affects offspring brain development, particularly myelination patterns and GABA receptor expression, with effects that persist into juvenile periods [43]. Additionally, the effects of single or repeated betamethasone exposure have been shown to affect HPA-axis function in both first and second-generation [52, 53]. These transgenerational effects show distinct sex-specific patterns and suggest that clinical use of antenatal steroids may have longer-lasting consequences than previously recognized.

Ethanol exposure models were also well-characterized, providing important insights into fetal alcohol spectrum disorders. These studies typically utilized chronic exposure protocols, with ethanol administered either through drinking water or direct administration [27, 33]. Key findings demonstrated that prenatal ethanol exposure affected multiple systems, including altered HPA axis function, modified glutamate signaling, and immune system development, particularly alveolar macrophage function [57].

Several other pharmacological interventions were studied and provided insights into programming effects of modulating the maternal HPA axis. Vartazarmian et al., [65] investigated prenatal selective serotonin reuptake inhibitor (fluoxetine) exposure and demonstrated altered pain thresholds in adult offspring despite no effects on pregnancy outcomes while Kaiser et al., [72] examined the effects of adrenocorticotropic hormone administration during pregnancy, finding increased aggressive behaviors and elevated cortisol levels in female offspring.

### Physiological models

Physiological models focused primarily on three key areas: growth restriction (uterine artery ligation), hypoxia exposure, and infection/inflammation challenges. Growth restriction models produced fetal growth restriction while maintaining pregnancy viability, allowing investigation of both immediate and long-term consequences of restricted fetal growth [29, 30, 54]. Cumberland et al., [45] demonstrated that the combination of growth restriction and prenatal stress increased the risk of preterm labor, while growth restriction alone did not affect gestation length. These models have also shown impacts on fetal endocrine function, with growth restriction combined with prenatal stress resulting in decreased fetal circulating cortisol levels and altered allopregnanolone concentrations. Some studies have explored potential therapeutic interventions, such as N-acetylcysteine supplementation, showing promising results in ameliorating some adverse effects of growth restriction, particularly regarding vascular function and oxidative stress markers [54, 56].

Nutritional models have demonstrated diverse impacts of maternal dietary deficiencies and restrictions on fetal/offspring development. Models specifically targeting a particular nutrient deprivation (maternal vitamin C or iron deficiencies) indicated particular gestational or postnatal age effects. For example, maternal vitamin C deficiency caused transient intrauterine growth restriction at gestational day 45 which resolved by gestational day 56 [61]. Iron deficiency implemented from pre-conception through lactation resulted in elevated cortisol in offspring at postnatal day 24 that normalized by day 84, suggesting developmental programming effects on stress responsiveness that were correctable with postnatal nutritional rehabilitation [29]. Global nutritional restrictions provided insights into placental adaptations and fetal growth programming including placental transcriptome changes relating to DNA repair, apoptosis regulation, nutrient transport and IGF1 regulation [30] and sex-specific effects on hypoxia and oxidative stress markers in the fetal brain [31]. Collectively, these nutritional models demonstrate that different types and severities of maternal dietary restrictions produce distinct patterns of fetal adaptation, from transient growth effects with micronutrient deficiencies to more profound metabolic and transcriptional changes with global caloric restriction.

Infection and inflammation models provided crucial insights into mechanisms of preterm birth and fetal inflammatory responses. Studies using various pathogens, including E. coli [63], Brucella [64], and cytomegalovirus [66], have increased understanding of maternal-fetal transmission and fetal immune responses. These models showed that timing and severity of infection significantly impact outcomes, with early gestation infections often leading to pregnancy loss [64, 66], while later infections resulted in fetal inflammation without pregnancy termination [63]. The work by Hensel et al., [64] particularly demonstrated how maternal infection can lead to placental inflammation and fetal compromise, with specific patterns of inflammatory markers that parallel human responses.

Hypoxia models were used to understand fetal adaptations to reduced oxygen availability, with protocols ranging from chronic moderate hypoxia to more severe intermittent exposures and carbon monoxide exposure [48, 50, 55, 56, 59]. These studies revealed important insights into fetal cardiovascular adaptations, including changes in endothelial nitric oxide synthase expression and vascular remodeling, and redox imbalances in the neonatal brain. Fetal organs respond differently to hypoxic challenges, with tissue-specific alterations in oxidative stress markers and cellular adaptation mechanisms [59]. Whilst studies that extended the understanding of hypoxia effects beyond the fetal period and demonstrated long-term cardiovascular programming in female offspring [55] and mechanistic understanding of how prenatal hypoxia may predispose neuronal dysfunction in adulthood [50].

## Discussion

This systematic review aimed to evaluate and synthesize the available literature on maternal stress during pregnancy using translational guinea pig models: Summarized in Figure 2. Overall, analysis of the available literature revealed a predominant focus on fetal and offspring outcomes, with relatively limited examination of maternal physiological adaptations to various stressors. While experimental designs successfully demonstrated diverse methods of increasing maternal stress, from acute psychosocial stressors to chronic physiological challenges, maternal responses were often only superficially characterized through basic measures such as weight gain and occasional cortisol sampling. Even when maternal stress was confirmed through elevated cortisol levels or behavioral changes, the underlying maternal adaptations and potential compensatory mechanisms remained largely unexplored. This knowledge gap is particularly notable given that maternal physiological responses likely play a crucial role in mediating the well-documented offspring effects, which frequently showed sex-specific patterns in behavioral, endocrine, and developmental outcomes.

**Figure 2.**
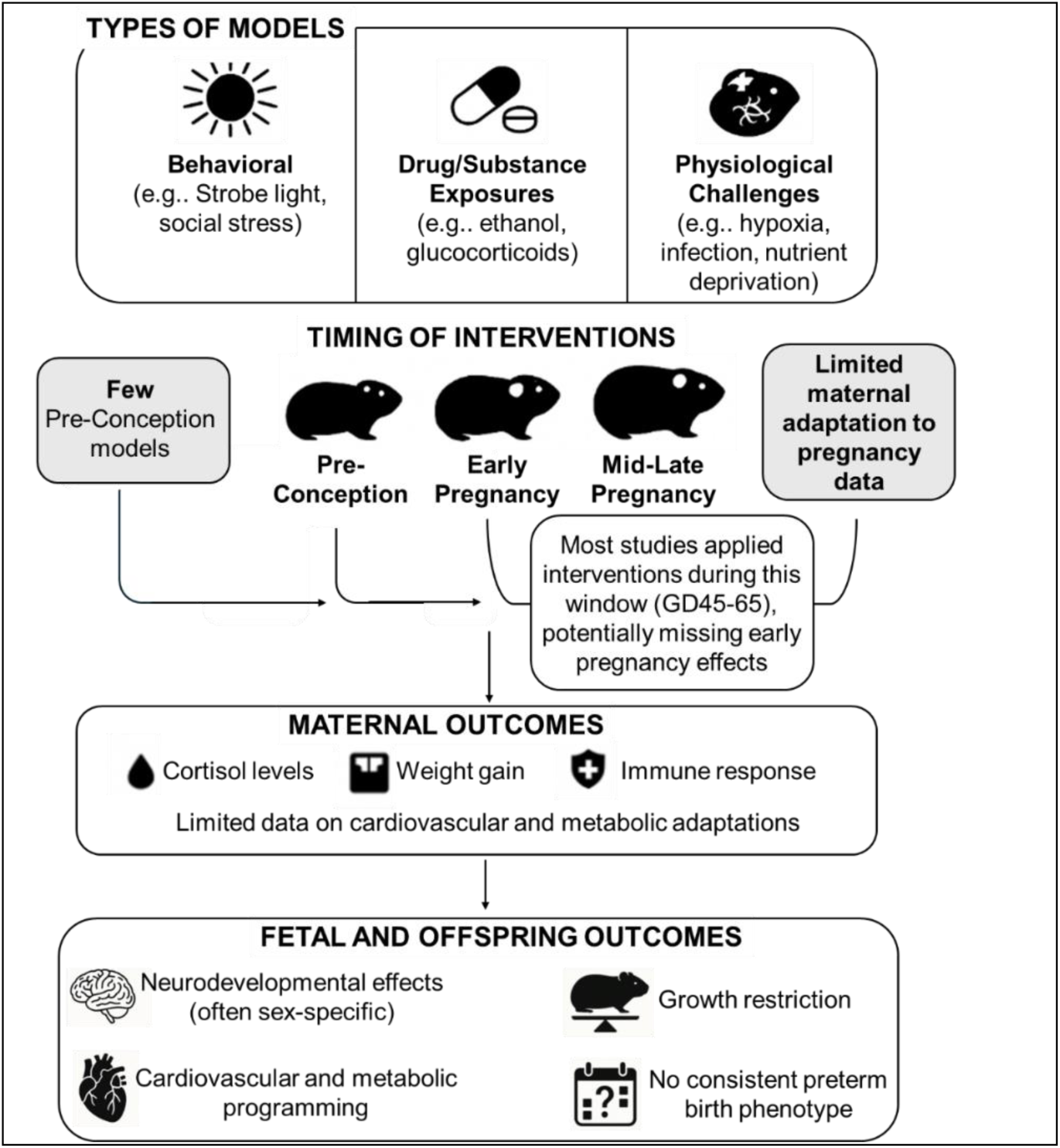
Summary of systematic review of the literature aimed to evaluate and synthesize the available literature on maternal stress during pregnancy using translational guinea pig models.

Of the studies that examined maternal responses, most focused on basic physiological markers such as weight gain and stress hormones. Maternal weight gain was generally maintained during stress interventions [34, 40, 41, 62], with reductions only observed in specific challenges like iron deficiency [29] and methadone treatment [47]. These findings parallel human studies where severe nutritional deficiencies or substance use affect maternal health [73, 74]. Maternal HPA axis activation was primarily assessed through cortisol measurements in blood or saliva. Elevated cortisol levels were observed following acute stressors such as strobe light exposure [37, 45] and combined environmental stressors [34]. Interestingly, several interventions including chronic ethanol exposure [27, 33] and iron deficiency [29] did not elicit increased cortisol responses, suggesting a possible adaptation of the maternal stress response system to chronic stress. Maternal adaptations to chronic maternal stress have also been observed in human pregnancies with chronic stress exposure [75]. These adaptations include changes in the HPA-axis function and cortisol patterns that attenuate stress reactivity and blood pressure responses [76]. Cardiovascular and metabolic adaptations have also been studied revealing differential hemodynamic responses and neuroendocrine reactivity [75]. Based on the literature reviewed here, the most severe maternal outcomes were observed in infection models [64, 66], particularly with high pathogen doses, mirroring the significant maternal morbidity associated with severe infections during human pregnancy [77, 78]. However, detailed characterization of maternal immune responses, cardiovascular adaptations, metabolic changes and behavioral modifications were largely absent from these studies. Overall leaving critical gaps in our understanding of how maternal systems adapt to maintain pregnancy despite significant stressors.

Despite extensive evidence linking maternal stress to preterm birth in human populations, the experimental models reviewed here showed little impact on gestational length, even with significant maternal stress manipulations. Cumberland et al., [45] demonstrated increased preterm labor and specifically when intrauterine growth restriction was combined with prenatal stress, whilst Maki et al., [31] reported preterm delivery in 3 of 18 nutrient restricted dams. These finding suggest that multiple ’hits’ may be necessary to initiate mechanisms resulting in spontaneous preterm delivery and highlights a critical gap in our understanding of the mechanisms linking maternal stress to pregnancy termination. Studies identified in the literature search that did result in spontaneous preterm birth but were subsequently excluded because no maternal stress experimentation was utilized, were artificially induced through pharmacological means (aglepristone and oxytocin)[79–81]. Pharmacological induction of parturition was used as a tool to study prematurity effects on fetal and offspring development rather than to understand spontaneous preterm birth mechanisms. Even in infection models, which showed the highest rates of pregnancy loss [63, 64, 66], the focus remained primarily on fetal outcomes rather than examining the maternal inflammatory and endocrine pathways leading to pregnancy termination. This gap in mechanistic understanding may partly explain why current clinical interventions for preventing preterm birth remain largely ineffective [82, 83], as they target downstream processes rather than the fundamental response pathways that initiate premature labor.

On of the goals of this systematic review was to synthesize the available literature on maternal stress during pregnancy using guinea pig models because of the translational potential guinea pigs offer. Traditionally, rodent models (particularly rats and mice) have been extensively used to study pregnancy and parturition. However, their fundamental differences in maternal adaptation to pregnancy responses and parturition initiation mechanisms limit their translational relevance for understanding human pregnancy physiology. Unlike humans and guinea pigs, where progesterone remains elevated throughout gestation and labor occurs without systemic progesterone withdrawal [19, 84, 85], mice and rats require a sharp decline in circulating progesterone to initiate parturition [19]. This distinct endocrine profile means that stress-induced or inflammation-mediated preterm birth in these species may operate through mechanisms that are not necessarily relevant to human pregnancy. Additionally, humans and guinea pigs shift progesterone production from the ovary to the placenta during pregnancy [86] so that pregnancy maintenance is independent of the ovary [87]; this mechanism is not observed in mice and rats [88]. Placental establishment and micro-structure in the guinea pigs more closely mirrors humans than mice and rats and guinea pigs deliver precocial young making which offers advantages in studying fetal and offspring responses to various experimental approaches. Maternal stress and glucocorticoid responses are also similar between humans and guinea pigs with cortisol, as opposed to corticosterone in mice and rats, being the primary glucocorticoid [22]. However, despite the similarities between humans and guinea pigs, the studies reviewed here demonstrate that even guinea pig models have not fully replicated the stress-induced preterm birth phenomenon observed in human populations. This suggests that additional factors, perhaps related to the complexity of human psychosocial stress or the interaction of multiple physiological systems, may be necessary to fully model the pathways leading to stress-induced preterm birth.

Another limitation in guinea pig models included in this review is that experimental approaches often began during established pregnancy rather than before conception or during early pregnancy. In humans, many risk factors for adverse pregnancy outcome are present before pregnancy, including chronic medical conditions (diabetes, hypertension), previous reproductive history, and social determinants of health (poverty, stress, racism) [89–92]. Human studies demonstrate that chronic maternal stress and inflammation present before conception significantly increase preterm birth risk, with one systematic review showing a 2-3 fold increased risk in women with pre-pregnancy anxiety or depression [93]. For many of the studies presented here, various maternal insults - including stress [36, 41], infection [63, 64], growth restriction [54], and drug exposures [27, 33] - created significant fetal effects, however, maternal physiology and health outcomes were often not reported. It is possible that the lack of reports on maternal health outcomes was because there was no significant effect to report. Hence furthering our hypothesis that experimental protocols that begin during pregnancy fail to capture the complex interplay of pre-pregnancy and early pregnancy factors that epidemiological studies have shown contribute to adverse pregnancy outcomes in humans [94, 95]. It is our belief that to develop more clinically relevant models, experimental manipulations likely need to begin prior to conception to better reflect human conditions where pre-existing maternal conditions or early pregnancy events contribute to adverse pregnancy outcome risk. This represents an important gap in the current literature and an opportunity for future research directions that could better align animal models with human clinical presentations.

## Author Contributions

RLW: Conceptualization, Methodology, Validation, Investigation, Formal Analysis and Writing. CM: Methodology, Validation, Investigation, Editing. AFL-J Methodology, Validation, Investigation, Editing. All authors approve final manuscript.

